# Daptomycin resistance in *Enterococcus faecium* can be delayed by disruption of the LiaFSR stress response pathway

**DOI:** 10.1101/2020.06.23.168401

**Authors:** Amy G. Prater, Heer H. Mehta, Kathryn Beabout, Adeline Supandy, William R. Miller, Truc T. Tran, Cesar A. Arias, Yousif Shamoo

**Author notes:** Corresponding author: Yousif Shamoo, Department of BioSciences, Rice University, 6100 Main St., Houston, TX.

## Abstract

The LiaFSR signaling pathway plays a major role in mediating daptomycin (DAP) resistance for both *Enterococcus faecalis* and *Enterococcus faecium*. LiaFSR inhibition induces DAP hypersusceptibility but could also potentially delay the acquisition of DAP resistance in a combinatorial therapy of DAP with a LiaFSR inhibitor. To evaluate the potential efficacy of this approach, the adaptation to DAP by both *E. faecalis* and *E. faecium* lacking a functional LiaFSR were examined. Here, clinical isolates of *E. faecium* with *liaR* deletions were evolved to DAP resistance using *in vitro* experimental evolution. Genomic analysis of resistant populations was used to identify both the alleles and their relative frequencies in driving DAP resistance. Microscopic and biochemical analyses were then employed to investigate how those adaptive alleles contributed to DAP resistance. We found that deletion of *liaR* from the *E. faecium* genome significantly delayed the onset of DAP resistance. Unsurprisingly, resistance strategies emerged eventually. These alternative strategies were influenced by both environment and ancestral genome. The delay in the acquisition of DAP resistance when *liaR* was deleted supports the concept of developing a LiaFSR pathway inhibitor to prolong DAP efficacy against enterococci. The loss of a functional LiaFSR pathway reset the adaptive landscape and forced adaptation to progress in new ways that were slower in providing DAP tolerance. The observed adaptive trajectories were strongly influenced by both the environment and ancestral genome.

## Introduction

Daptomycin (DAP) is a cyclic lipopeptide antibiotic used to treat multi-drug-resistant Gram-positive infections (1–3). The DAP mechanism remains under investigation but is known to form a positively charged complex in the presence of calcium and binds to phosphatidylglycerol-rich regions of the membrane. The DAP:Calcium complex inserts into the membrane, interacts with lipid II precursors causing inner protein mislocalization which then leads to cell death (4, 5). Specifically, recent work in *Staphylococcus aureus* has suggested that DAP-mediated killing is the result of the DAP:Ca^+2^ complex binding to undecaprenyl-coupled cell envelope precursors in the presence of phosphatidylglycerol (6).

Unfortunately, DAP resistance is increasingly observed in the clinic and commonly attributed to one of two main mechanisms: repulsion of the antibiotic from the cell surface or redistribution of the antibiotic from critical septal sites. Repulsion is often seen in *S. aureus* and is characterized by an increase in the net surface charge. This is regularly mediated by the incorporation of positively charged residues into either lipoteichoic acids (LTAs) or phosphatidylglycerol through changes in the *dltABCD* pathway or *mprF*, respectively (7–9). Redistribution is commonly observed in *Enterococcus faecalis* where activating mutations in *liaFSR* and cardiolipin synthase (*cls*) redistribute anionic cardiolipin microdomains away from the division septa to possibly prevent DAP from affecting the cellular machinery required for efficient septation (5, 10, 11).

While the redistribution phenotype is not typically observed in DAP-resistant *E. faecium* (12), *liaFSR* mutations are commonly present and contribute to DAP tolerance (13). The occurrence of these mutations across enterococcal species reveals a potential target for the generation of an “anti-evolution” co-drug. This potential therapy could be administered alongside DAP to extend clinical DAP efficacy by inducing hypersusceptibility, delaying the evolution of resistance, and reducing the spread of resistant strains (14).

Previously, we determined that additional mutations were necessary to achieve high levels of DAP resistance in DAP-tolerant *E. faecium* containing LiaFSR-activating alleles (12). Using *in vitro* experimental evolution within flask-transfers, *yvcRS* mutations were identified and correlated with increased *dltABCD* and *mprF* transcripts and DAP repulsion (12). Alternatively, in a biofilm-heavy bioreactor environment, a single nucleotide polymorphism (SNP) in *divIVA*, associated with increased DAP resistance, resulted in severe septal abnormalities. Because these additional resistance methods are accessible to *E. faecium*, it is possible that these same strategies could be readily available to *E. faecium* lacking a functional LiaFSR system, and that DAP resistance would be quickly achieved without LiaFSR activity-rendering a potential LiaFSR inhibitor less effective.

Here, we evolved two clinical *E. faecium* isolates, HOU503 and HOU515 (13), with the *liaR* response regulator deleted from their genomes (503FΔ*liaR* and 515FΔ*liaR*, respectively) (15), to DAP resistance using both flask-transfer and bioreactor-mediated experimental evolution, as described previously (12). These two methods promote different adaptive environments which can select for potentially different resistance strategies and allow for a more thorough survey of evolutionary trajectories leading to resistance. Using these *in vitro* techniques, we found that deletion of *liaR* greatly delayed the onset of DAP resistance, but that resistance was achieved eventually by both expected and unexpected means.

## Results

### Deletion of *liaR* significantly delayed the onset of DAP resistance

Previous experimental evolution studies in *E. faecium* (12) led to the prediction that alternative evolutionary trajectories could still be expected to provide DAP resistance when the LiaFSR signaling response was absent (12). The central question was whether these additional paths could readily compensate for the loss of LiaFSR and thereby rapidly confer DAP resistance. To test this hypothesis, 503FΔ*liaR* and 515FΔ*liaR* were evolved to DAP resistance in both flasks and a bioreactor. Five independent 503FΔ*liaR* and 515FΔ*liaR* populations each were evolved via flask-transfer, favoring planktonic populations as cells that adhere to surfaces are less likely to be transferred each day. Cells were transferred to increasing DAP concentrations until populations were growing at ≥8 mg/L DAP, resistant by CLSI standards (16) (Figure 1). 503FΔ*liaR* flask populations reached completion within 18 days while the 515FΔ*liaR* flask populations reached completion in 20 days. This timeline is significantly delayed compared to the six days required for the clinical HOU503 (12) and HOU515 flask populations to achieve DAP resistance.

**Figure 1.**
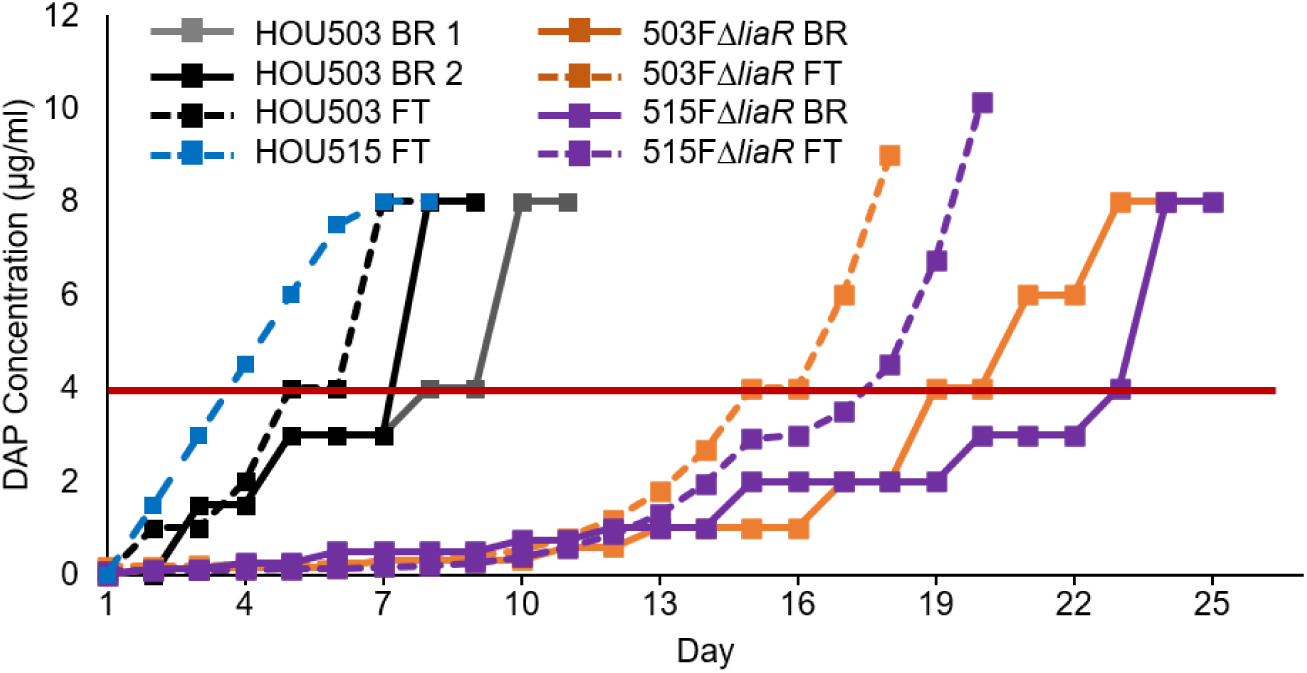
Deleting *liaR* delayed DAP resistance. 503FΔ*liaR* and 515FΔ*liaR* were adapted to DAP resistance via flask-transfer (FT; five populations each) or bioreactor (BR; one population each). The horizontal red line indicates the clinical DAP susceptibility cutoff. Each replicate flask-transfer population was transferred to identical DAP concentrations. HOU503 data are replotted from Prater *et al.*, 2019 for comparison.

503FΔ*liaR* and 515FΔ*liaR* were also evolved to DAP resistance in singlicate using a bioreactor, which selects for the formation of complex-structured communities that are typically found in biofilms (10, 12, 17). Bioreactor experiments maintain highly polymorphic populations, however, that polymorphism is also highly replicable when the selection is moderate and the population sufficiently large; previously, replicate bioreactor populations evolved similar mutations, though the frequency at which those mutations occurred can vary (18–21). The bioreactor was used to investigate whether the opportunity to form biofilms altered the observed evolutionary trajectories compared to the flask-transfer studies, as was observed in HOU503 (12). The 503FΔ*liaR* population reached 8 mg/L DAP within 23 days, compared to the eight or 10 days required for the parent, HOU503, while 515FΔ*liaR* reached completion in 25 days (Figure 1). In the absence of a functional LiaFSR system, the timeline to DAP resistance doubled in *E. faecium* regardless of adaptive environment

Below we report on the alternative, but common genomic resistance strategies, while less common adaptive trajectories, hitchhiking mutations, and plasmid-associated mutations can be found in the Supplemental Text, Figures S3-4, and Table S1. Of note, end-point isolates from the bioreactor studies were not randomly selected, but rather selected to sample a range of MICs, and thus the frequency of end-point genotypes does not reflect the frequency of those genotypes within the bioreactor.

### Repulsion-based DAP resistance evolved through changes in *yvcRS* signaling

Two 503FΔ*liaR* and all five 515FΔ*liaR* flask populations contained mutations in *yvcRS* (Tables 1-2), the multi-component system that senses bacitracin and was shown previously to provide DAP resistance in flask-transfer isolates of *E. faecium* HOU503 (12). In HOU503, *yvcRS* mutations were correlated with an increase in *dltABCD* transcripts, an increase in cell surface charge, and reduced DAP binding-consistent with the repulsion-based resistance mechanism (12). Here, we again found that mutations in *yvcRS* occurred, albeit without an intact LiaFSR system.

**Table 1.** 503FΔ*liaR* flask-transfer end-point isolate genotypes.

| Isolate | DAP MIC | Population | VAN Plasmid | *cls* | *yvcS* | *pyre* | *entfae_548* | *nraA* | *entfae_1207* | *rpoC* | *entfae_357* | *entfae_370* | *entfae_933* | *entfae_1482* | *entfae_2177* | *entfae_2936* |
| --- | --- | --- | --- | --- | --- | --- | --- | --- | --- | --- | --- | --- | --- | --- | --- | --- |
| 503FΔ*liaR* FT 1-1 | 32 | 1 | $\Delta$ | A20D | A647P | | | | | | | | | | | |
| 503FΔ*liaR* FT 1-2 | 16 | 1 | $\Delta$ | A20D | A647P | | | | | | | | N73N | | | |
| 503FΔ*liaR* FT 2-1 | 8 | 2 | $\Delta$ | R218Q | | +309 | T227R | | | | | | | | | |
| 503FΔ*liaR* FT 2-2 | 8 | 2 | $\Delta$ | R218Q | | +309 | T227R | | | | D108Y | | | | | |
| 503FΔ*liaR* FT 3-1 | >64 | 3 | $\Delta$ | A20D | | | | | | | | | | A39S | L303S | T299M |
| 503FΔ*liaR* FT 4-1 | 32 | 4 | $\Delta$ | | | | | H105P | Q387K | | | | | | | |
| 503FΔ*liaR* FT 4-2 | 32 | 4 | $\Delta$ | | | | | H105P | Q387K | | | | | | | |
| 503FΔ*liaR* FT 5-1 | 4 | 5 | | R211L | W503S | | | | | T777K | | | | | | |
| 503FΔ*liaR* FT 5-2 | 32 | 5 | | R211L | W503S | | | | | T777K | | L69L | | | | |
| Total Strains with Changes |  |  | 7 | 7 | 4 | 2 | 2 | 2 | 2 | 2 | 1 | 1 | 1 | 1 | 1 | 1 |

**Table 2.**
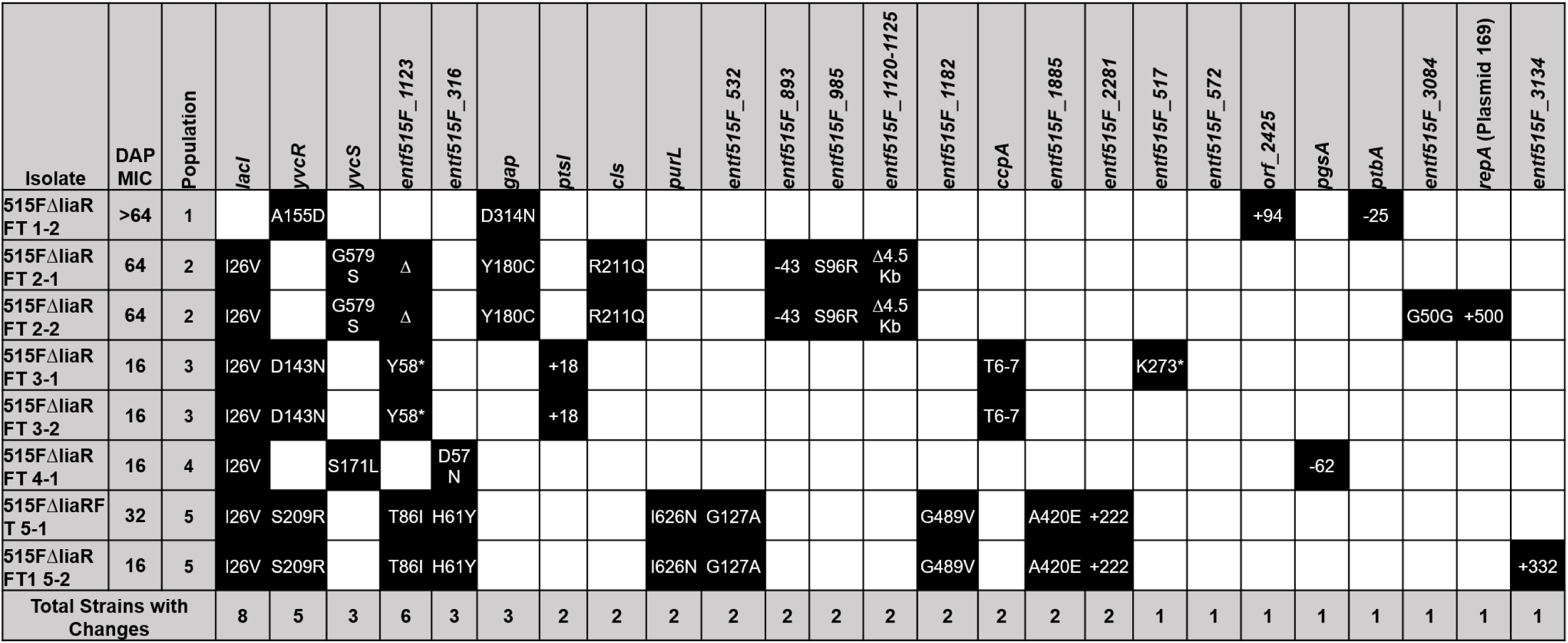
515FΔ*liaR* flask-transfer end-point isolate genotypes.

As was found in HOU503 flask-adapted isolates with *yvcRS* mutations (12), *dltA* transcripts were elevated compared to the housekeeping gene, glucose-1-dehydrogenase 4 (*gdhIV*), in Δ*liaR* flask-transfer isolates containing *yvcRS* mutations (Figure 2). Of note, without allelic replacements, the most parsimonious arguments between genotype and phenotype are shown here, where multiple isolates carrying similar mutations were assessed.

**Figure 2.**
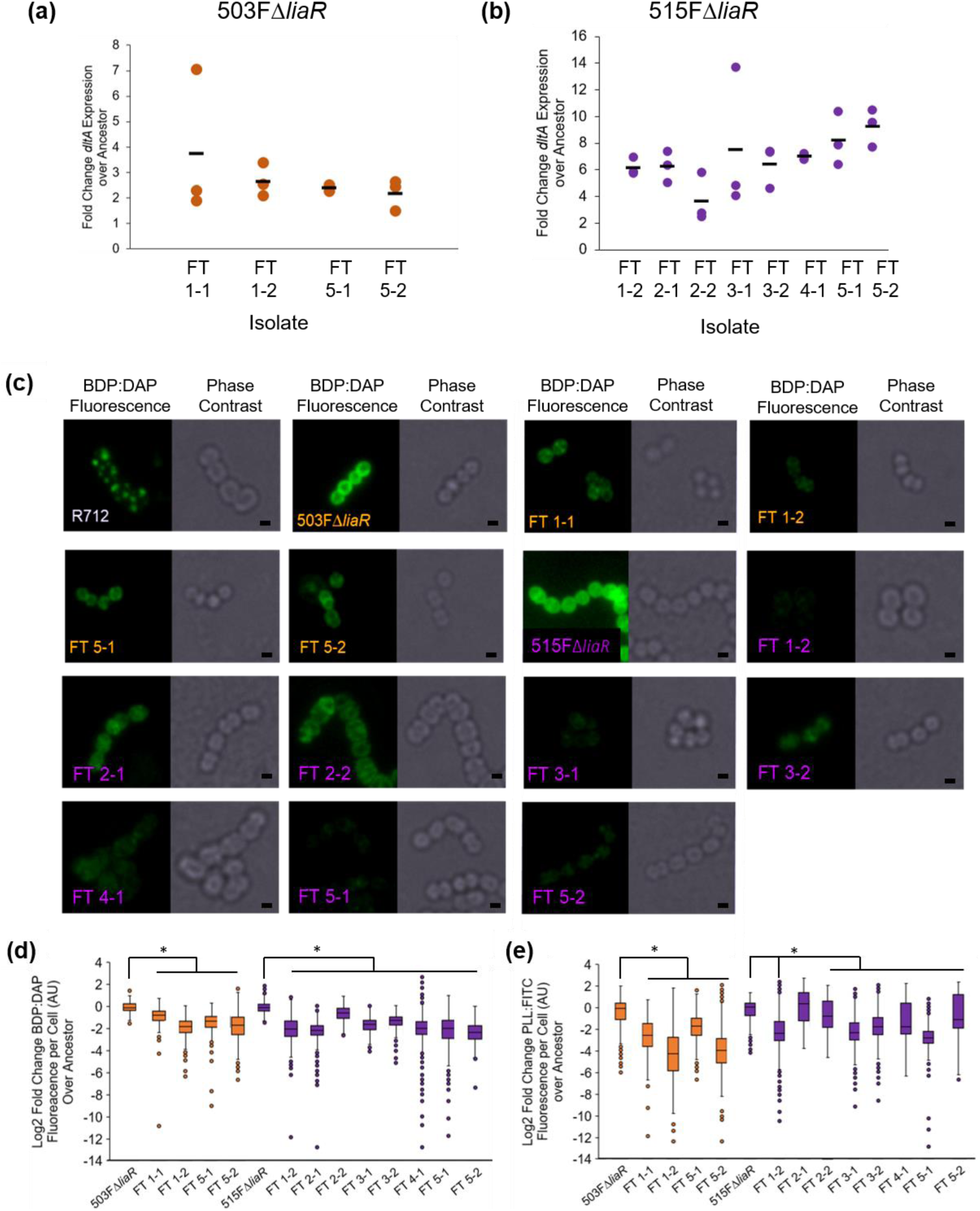
Flask-transfer isolates with mutations in *yvcRS* repulsed DAP. **(a)** qPCR of dltA transcripts for 503FΔliaR isolates using gdhIV as reference. **(b)** qPCR of dltA transcripts for 515FΔliaR isolates using gdhIV as reference. Circles indicate biological replicates while bars indicate mean. **(c)** Isolates were incubated with 32 µg/ml BDP-DAP. Scale bars show 1 µm. E. faecalis R712 acts as a control showing the DAP redistribution phenotype. (d) Quantification of BDP:DAP. **(e)** Quantification of PLL:FITC; images found in **Figure S1**. Isolate names in orange are 503FΔliaR isolates and names in purple are 515FΔliaR isolates. An asterisk indicates significance compared to the ancestor (p<0.05) using Mann-Whitney with post hoc Holm-Bonferroni adjustment. Experiment performed in duplicate on separate days. Quantification using ImageJ.

To determine cell surface charge, isolates were incubated with Poly-L-Lysine conjugated to FITC (PLL:FITC) and cellular fluorescence was quantified using fluorescence microscopy and ImageJ. If cells bind less PLL:FITC, it suggests an increase in cell surface charge consistent with repulsion-based DAP resistance (12, 22). All flask-transfer isolates containing *yvcRS* mutations bound less PLL:FITC than the ancestors (except 515FΔ*liaR* FT 1-2), indicating an increase in cell surface charge (Figures 2, S1). Similarly, isolates were incubated with DAP conjugated to the fluorophore bodipy (BDP:DAP) to determine DAP binding patterns (12, 13, 15, 23). Consistent with the increase in surface charge, flask-transfer isolates containing *yvcRS* mutations bound significantly less BDP:DAP than the ancestor (Figure 2). Incubation with 10-nonyl-acride orange (NAO) (5, 11, 24) revealed no evidence of lipid redistribution (Figure S4). Importantly, 503FΔ*liaR* FT1-1 (*yvcS*^A647P^, *cls*^A20D^) had similar mutations to HOU503 FT5 (*yvcS*^S23I^, *cls*^R218Q^)(12), yet the timelines to resistance differed markedly.

### Bioreactor-derived 503FΔ*liaR* isolate with *divIVA*^I92F^ resisted DAP largely through an increased surface charge

DivIVA is a scaffold protein at the division septum and cellular poles that aids in septum formation and chromosomal segregation (25). In a previous study, *divIVA*^*Q75K*^ in *E. faecium* HOU503 was found to be correlated with DAP resistance through complex mechanisms that remain under study (12). When *divIVA*^*Q75K*^ was present, DAP resistance was mediated by a combination of modest increases in cell surface charge but we speculate that in light of recent work on the mechanism of action of DAP in *S. aureus* (6), mutations to DivIVA may mitigate mislocalization of this critical cell division protein induced by DAP effects on the cell membrane.

Here, *divIVA*^I92F^ was first observed on Day 5 and comprised 52% of the final 503FΔ*liaR* bioreactor population (Figure 3a) and found in two end-point isolates (Table 3). Both I92F and Q75K are in the predicted loop region between two N-terminal coiled coils, suggesting the importance of modifications in this region for DAP resistance. Using isolate 503FΔ*liaR* P25 (*divIVA*^I92F^, *cls*^R211L^, *entfae_126*^V30*^), we found that this isolate produced abnormal division septa, bound less PLL:FITC (indicating a more positive surface charge) and some evidence of a subpopulation that hyperaccumulated DAP without redistributing anionic phospholipid microdomains (Figures 4, S1-S2). This was consistent with what was observed previously in the HOU503 *divIVA*^Q75K^ DAP-resistant isolates (12). The exact mechanism remains under investigation. Importantly, while we observed a similar mutation that contributed to rapid acquisition of high levels of DAP resistance in HOU503 (12), here, DAP adaptation in 503FΔ*liaR* was significantly delayed.

**Table 3.**
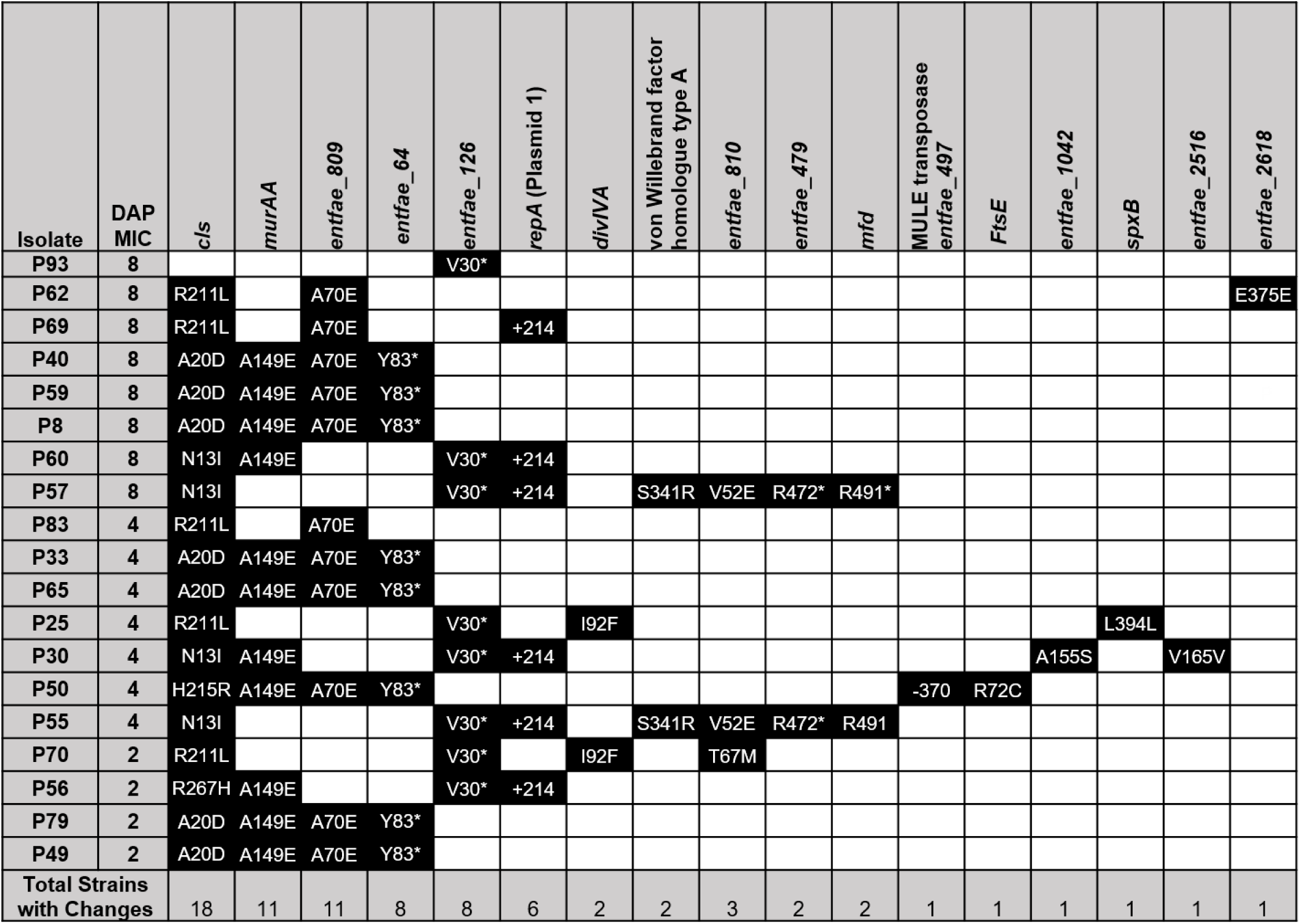
503FΔ*liaR* bioreactor-derived end-point isolate genotypes.

**Figure 3.**
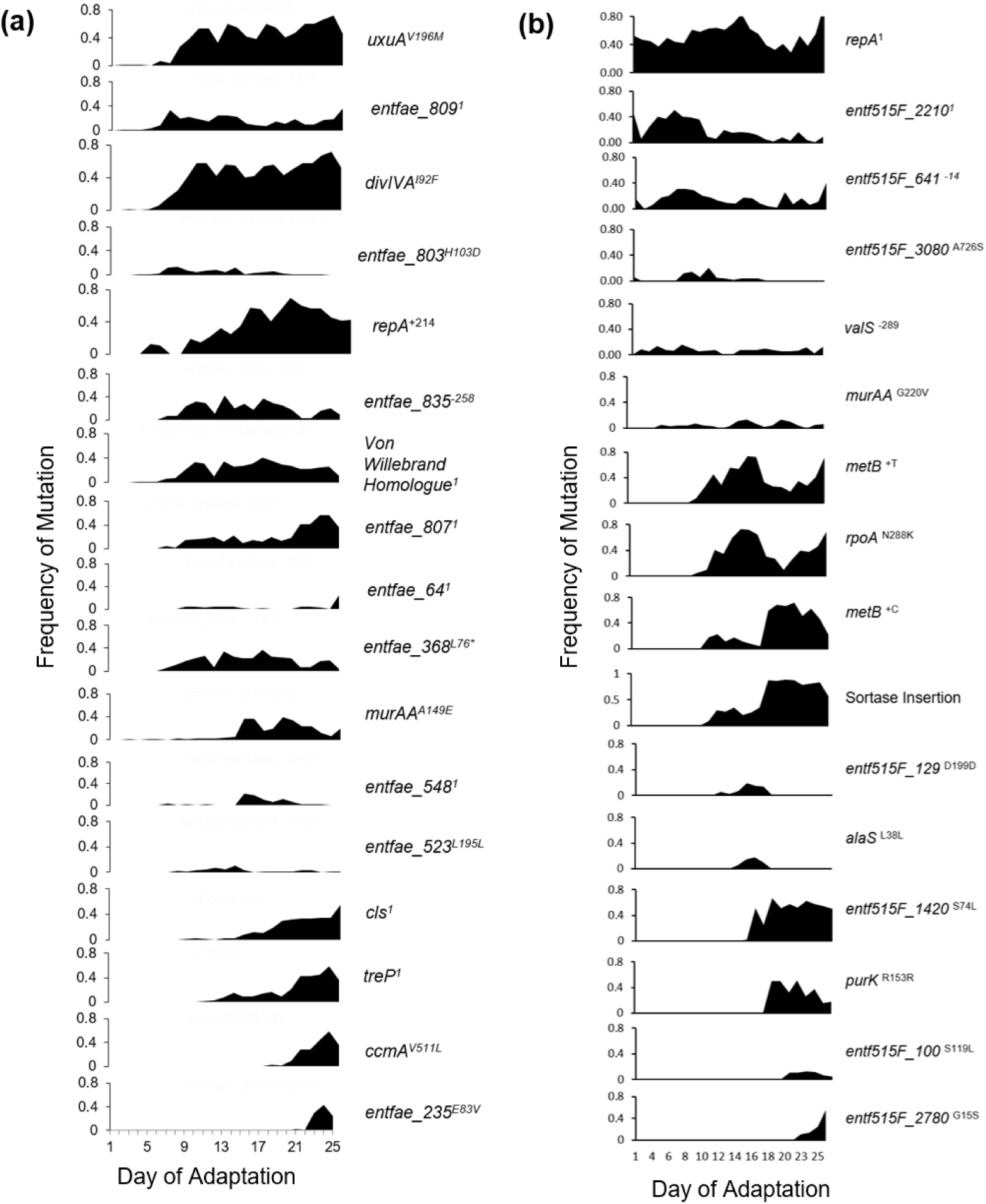
LiaR-independent bioreactor mutation frequencies over time. Mutations that reached a minimum of 10% frequency on two consecutive days are included here. 1denotes multiple mutations were present within that gene, the summation of which is represented here. **(a)** 503FΔliaR. **(b)** 515FΔliaR. Importantly, mutations unrelated to DAP resistance can accumulate within a population by “hitchhiking” with a bone fide adaptive mutation, such as uxuAV196M with divIVAI92F as seen in panel (a).

**Figure 4.**
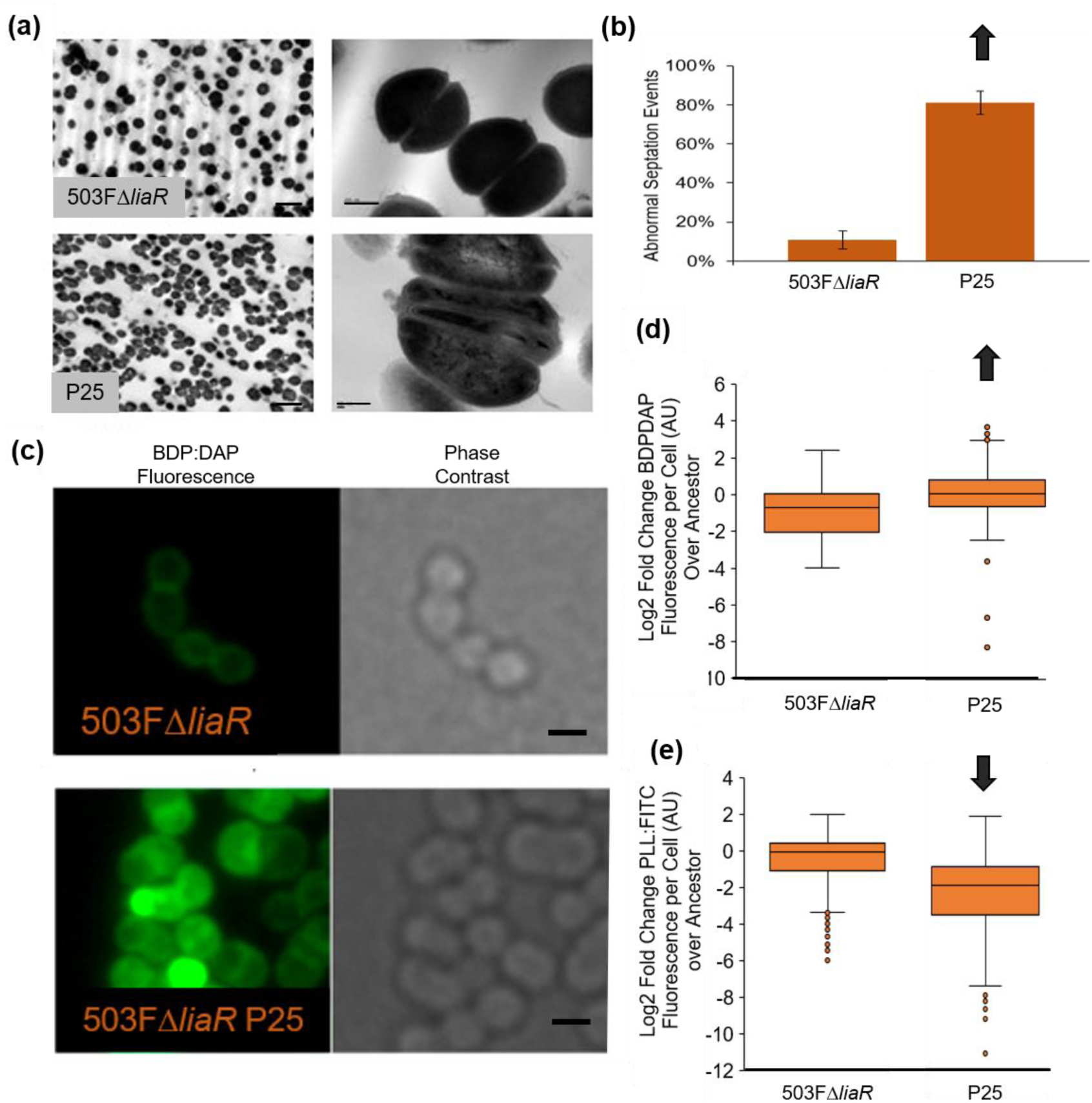
503FΔ*liaR* bioreactor-derived isolate with *divIVA*^I92F^ had aberrant septal formation. **(a)** TEM with scale bars indicating 2 µm (left) and 200 nm (right). **(b)** Aberrant septation events were quantified and significance determined by Student’s T test with (p<0.05). **(c)** Isolates incubated with 32 µg/ml BDP:DAP with scale bars indicating 1 µm. **(d)** Quantification of BDP:DAP using ImageJ. **E.** Quantification of PLL:FITC using ImageJ. Images found in **Figure S1.** Arrows indicate significance compared to the ancestor (p<0.05) using Mann-Whitney with *post hoc* Holm-Bonferroni adjustment. Experiment performed in duplicate on separate days.

### Bioreactor-derived 503FΔ*liaR* isolates with *murAA*^A149E^ may increase cell surface charge

MurAA catalyzes the first committed step in peptidoglycan synthesis by transferring enolpyruvate from phosphoenolphyruvate (PEP) to uridine diphospho-N-acetylglucosamine (UNAG). This reaction is targeted by the antibiotic fosfomycin (FOF), which irreversibly binds and inactivates the MurA active site (26, 27). The *murAA*^A149E^ allele was present at < 5% from Day 8-12 and grew to comprise 40% of the Day 18 population but then declined to 20% of the final population. *murAA*^A149E^ was present in 11/19 503FΔ*liaR* bioreactor-derived end-point isolates (Figure 3a and Table 3). These same isolates also exhibited increased FOF sensitivity with MICs falling from >32 to 8-16 mg/L. Interestingly, while MurAA is required for cephalosporin resistance (28), isolates containing *murAA*^A149E^ did not show a change in Cefotaxime MIC (remaining at 32 mg/L).

Two *murAA*^A149E^ isolates with the fewest mutations, 503FΔ*liaR* P8 (*murAA*^A149E^, *cls*^A20D^, *entfae_809*^A70E^, *entfae_64*^Y83*^) and 503FΔ*liaR* P60 (*murAA*^A149E^, *cls*^N13I^, *entfae_126*^V30*^), were selected for additional phenotypic analysis. Both isolates bound significantly less PLL:FITC than the ancestor without evidence of NAO remodeling. However, paradoxically, both isolates appear to bind more BDP:DAP than the ancestor (Figures 5, S3-S4). So, while the cells had a modest increase in cell surface charge, it was not correlated with DAP repulsion. The presence of secondary mutations makes it difficult to make a clear association of *murAA*^A149E^ to DAP resistance. However, the consistent identification of *murAA*^A149E^ has led us to begin biochemical and structural studies of MurAA and MurAA^A149E^. In general, isolates bound less PLL:FITC and did not redistribute lipid microdomains, suggesting that the changes in MurAA^A149E^ function may indeed contribute to DAP-tolerance by changing the physical properties of the cell wall.

**Figure 5.**
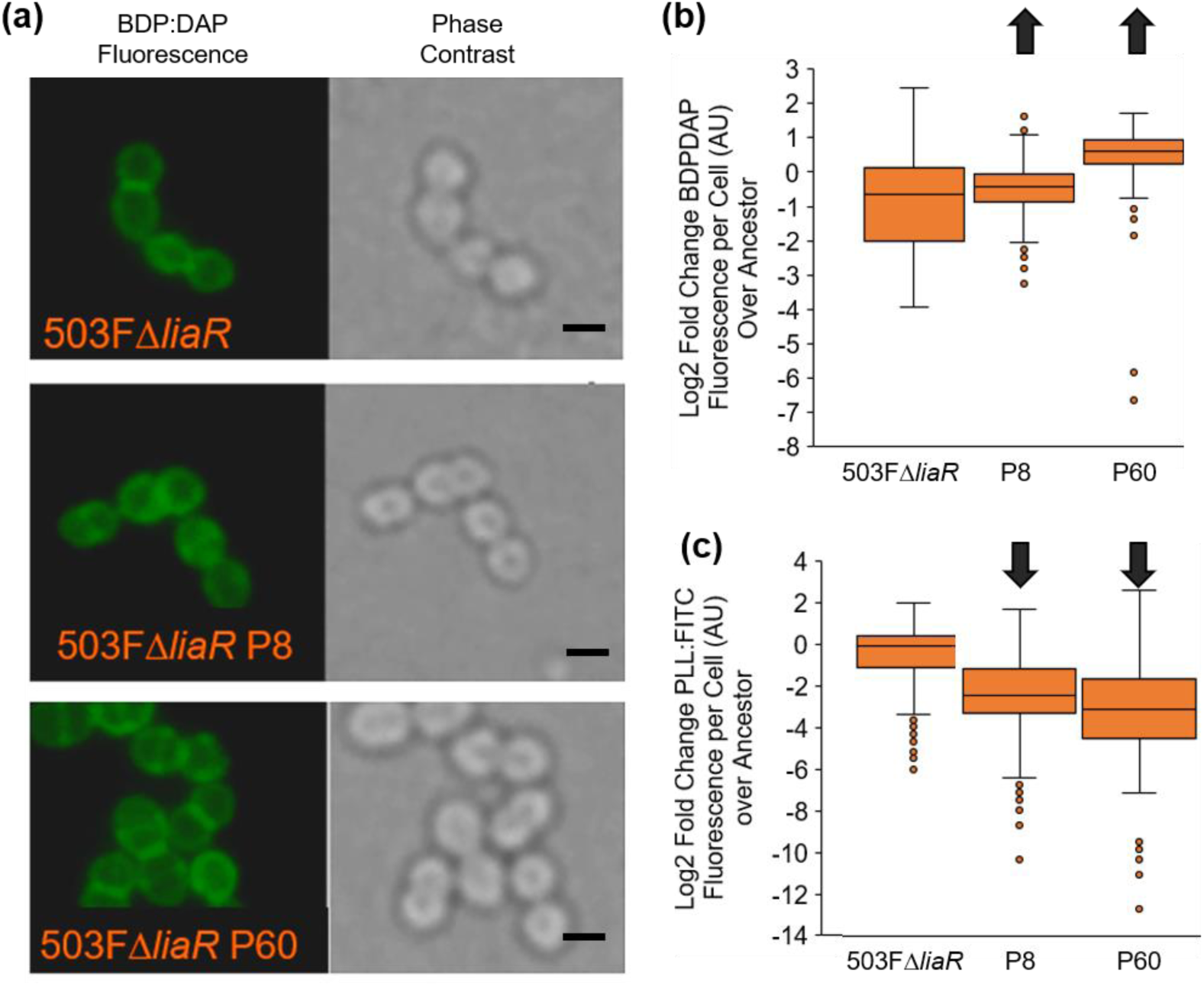
503FΔ*liaR* bioreactor-derived isolates with *murAA*^A149E^ may increase cell surface charge. **(a)** Isolates incubated with 32 µg/ml BDP:DAP. Scale bars indicate 1 µm. **(b)** Quantification of BDP:DAP using ImageJ. **(c**) Quantification of PLL:FITC using ImageJ. Images found in **Figure S1.** Arrows indicate significance compared to the ancestor (p<0.05) using Mann-Whitney with post hoc Holm-Bonferroni adjustment. Experiment performed in duplicate on separate days.

Interestingly, in 515FΔ*liaR, murAA*^G220V^ was observed in three bioreactor-derived end-point isolates, each of which had an additional 12-33 mutations (Table 4). An isolate containing this mutation (P53) also exhibited a minor increase in cell surface charge without evidence of DAP repulsion (Figure 6). This *murAA* mutation was located alongside nine additional mutations over 460 Kb that were identical to mutations acquired by a hypermutator subpopulation. It is likely that these mutations were part of a homologous recombination event. Further discussion can be found in the Supplemental Text.

**Table 4.**
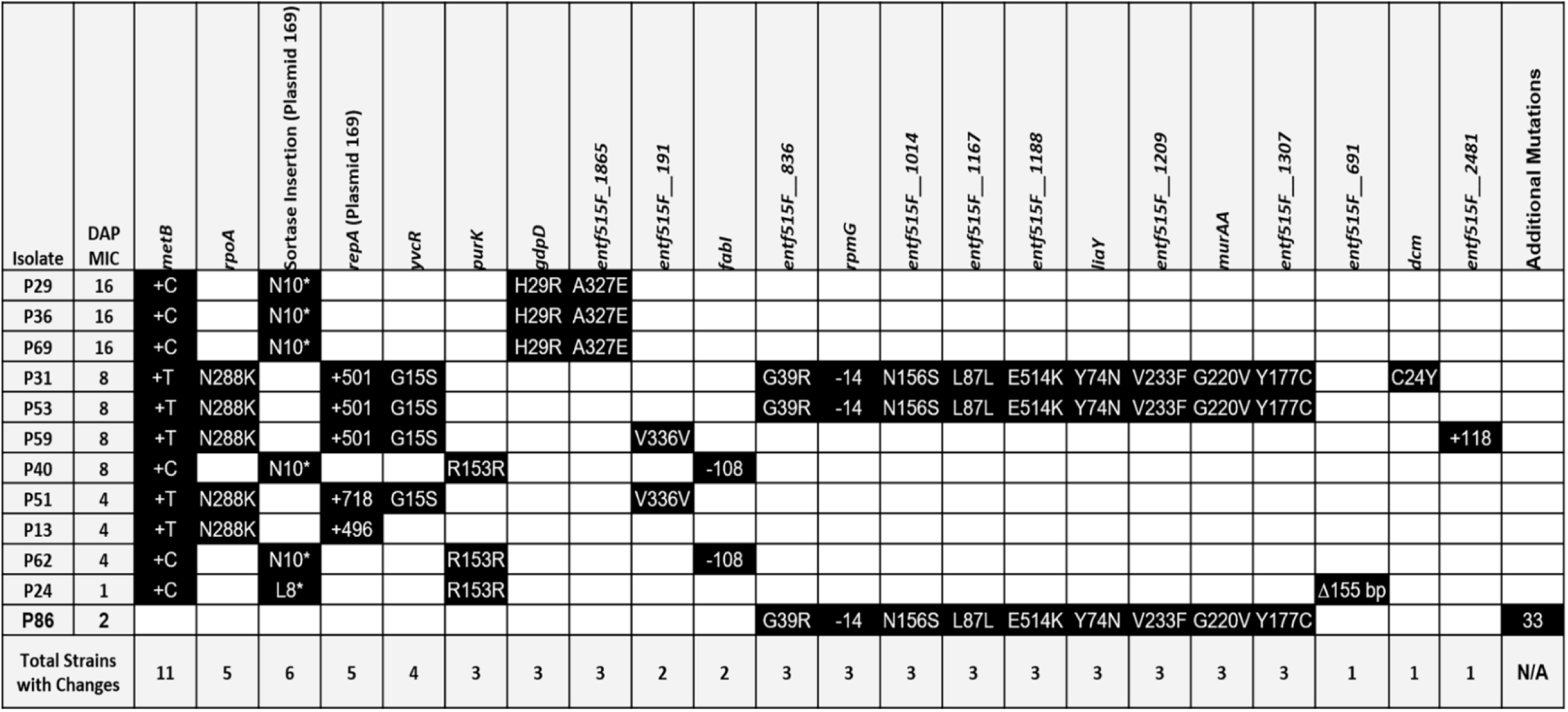
515FΔ*liaR* bioreactor-derived end-point isolate genotypes.

**Figure 6.**
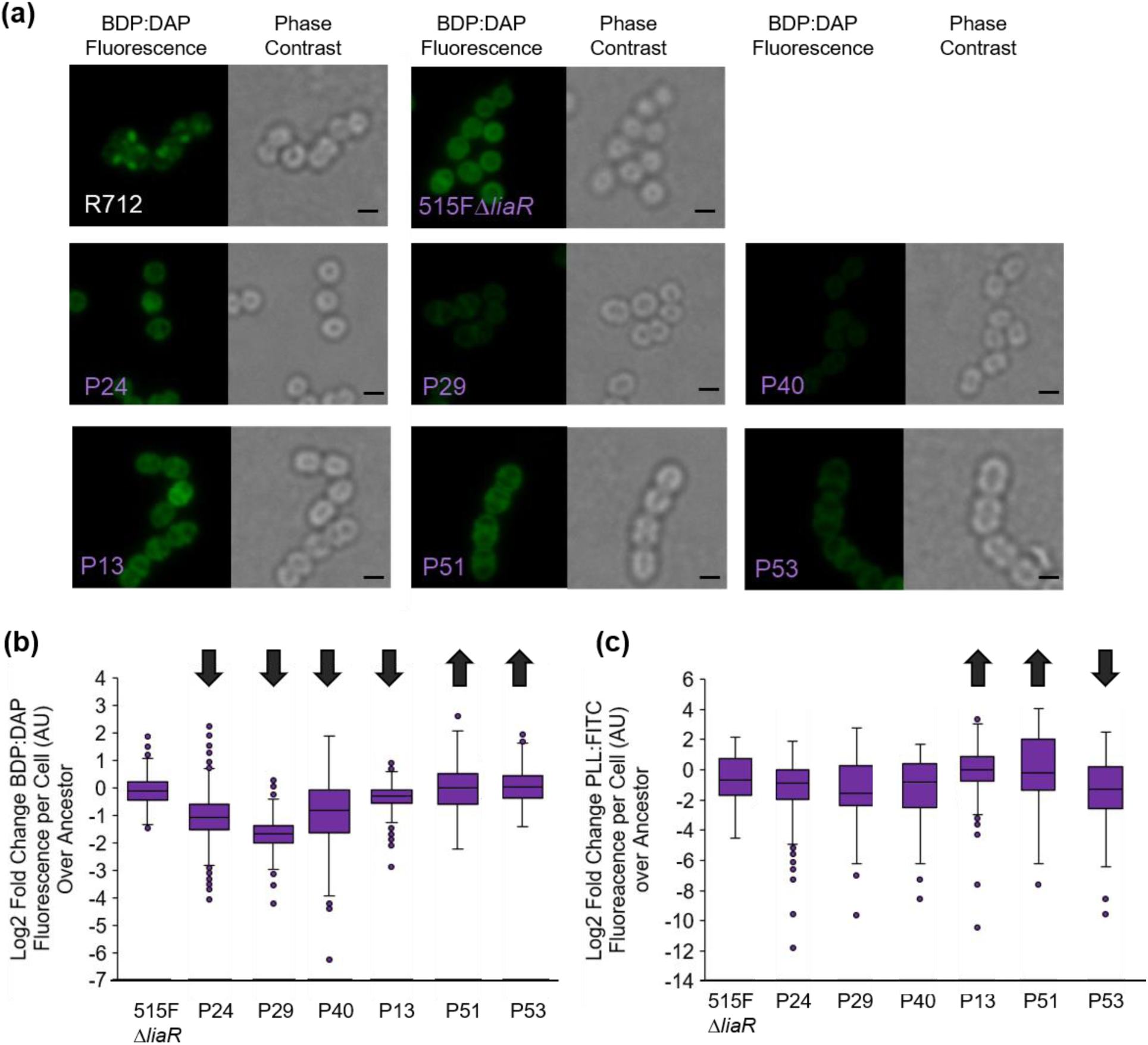
515FΔ*liaR* bioreactor-derived isolates with *metB* mutations produced variable phenotypes. **(a**) Isolates incubated with 32 µg/ml BDP:DAP. Scale bars indicate 1 µm. **(b)** Quantification of BDP:DAP using ImageJ. **(c)** Quantification of PLL:FITC using ImageJ. Images found in **Figure S1.** Arrows indicate significance compared to the ancestor (p<0.05) using Mann-Whitney with post hoc Holm-Bonferroni adjustment. Experiment performed in duplicate on separate days.

### *cls* mutations were commonly acquired

Mutations in *cls* are commonly found in DAP-resistant enterococci (10, 12, 13, 29, 30), however they are neither necessary nor sufficient to confer DAP resistance (31). Here, we found *cls* mutations in four 503FΔ*liaR* flask-transfer populations, one 515FΔ*liaR* flask-transfer population, and in 34% of the final 503FΔ*liaR* bioreactor population. Their presence here again indicates the importance of membrane modification, even without a functional LiaFSR system. Interestingly, all mutations clustered in the transmembrane or PLD1 domains, consistent with what has been observed previously (10, 12, 13, 29, 30), and indicating the importance of modification to these domains in DAP resistance.

### Mutations in *metB* were commonly acquired in bioreactor-evolved 515FΔ*liaR* resulting in complex phenotypes

In 515FΔ*liaR, entf515F_2516* was annotated as *metB*, which in *E. coli* encodes cystathionine gamma-synthase and catalyzes the formation of L-cystathionine and succinate from L-cysteine and O4-succinyl-L-homoserine (32). This reaction plays a role in sulfur, cysteine, and methionine metabolism. The *metB* mutations first appeared on day 9 and 10 and rose in frequency to comprise a combined 94% of the final 515FΔ*liaR* population and present in 11/12 end-point isolates (Figure 3 and Table 4). These mutations were characterized by a single base pair insertion of either cytosine or thymine after nucleotide 634. This insertion resulted in an extension of the open reading frame into the downstream gene, *entf515F_2515* (annotated as cys/met metabolism; PLP dependent enzyme family protein; Figure S6). Essentially, this mutation resulted in a fusion of these two gene products. In addition to this fusion, each *metB* lineage also contained a mutation affecting the sequence of Plasmid 169, either an insertional element into a Type A Sortase or a SNP downstream of *repA*. This fusion event and the possible effects of these plasmid-encoded mutations are discussed further in the Supplemental Text. Both *metB* mutations were acquired early in adaptation (prior to most ancillary mutations) and were not identified in a no-DAP control adaptation (Supplemental Text), indicating their potential significance in contributing to DAP resistance.

Generally, each *metB* lineage produced differing phenotypes, likely resulting from the differing secondary mutations present within each genome. Changes in *metB* are likely to have strongly pleiotropic effects on metabolic flux that could also increase DAP resistance and will require more investigation. The *metB*^+C^ lineage (P24, P29, P40) showed evidence of a potential non-charge based DAP repulsion mechanism (Figure 6 and S3), though more conclusive analysis using allelic replacements with *metB*^+C^ would be required to support this hypothesis. Conversely, *metB*^+T^ isolates (P13, P51, P53) showed more complex and varied phenotypes of differential PLL:FITC binding. P13 bound less BDP:DAP than the ancestor while P51/P53 bound slightly more BDP:DAP (Figure 6 and S3). Incubation with NAO, however, revealed a unique, patchy phenotype in *metB*^+T^ isolates that did not correspond to patchy BDP:DAP binding, suggesting lipid remodeling without DAP remodeling (Figure 7). Interestingly, Isolates P51 (*metB*^+T^, *rpoA*^+500^, *repA*^+718^, *yvcR*^G15S^, *entf515F_191*^V336V^) and P29 (*metB*^+C^, sortase insertion, *gdpD*^H29R^, *entf515f*_1865^A327E^) produced a unique phenotype where NAO staining condensed to a single focal point in cells, creating a dotted pattern (Figure 7). The lack of similar alleles between these two isolates suggests that this dot phenotype was the result of convergent evolution and is discussed further in the Supplemental Text. However, the lack of a clear phenotype between isolates containing differing *metB* mutations suggests that the ancillary mutations make key contributions to the DAP resistant phenotype through epistatic interactions.

**Figure 7.**
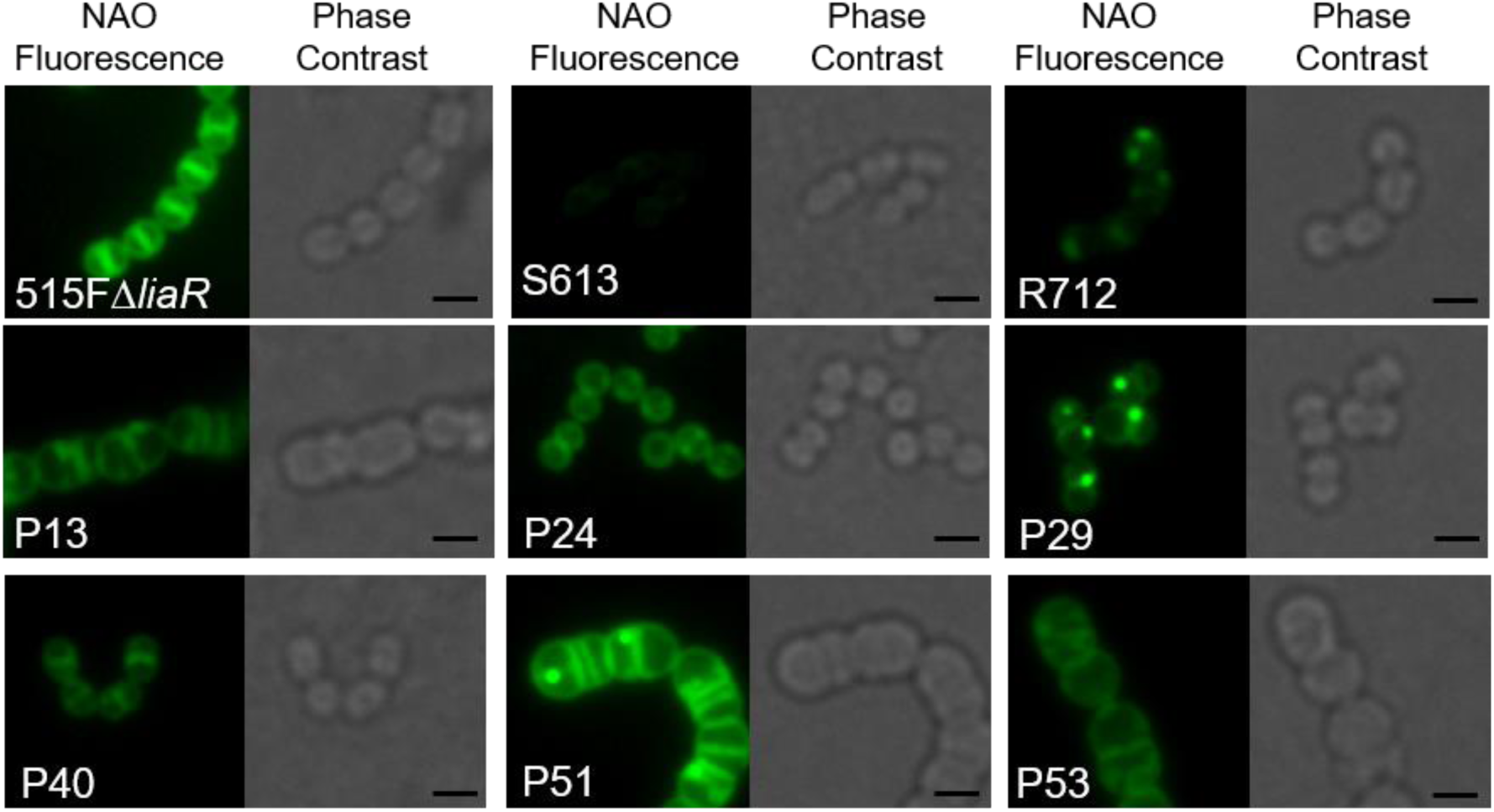
515FΔ*liaR* bioreactor-derived isolates with *metB* mutations produce interesting membrane structures. Isolates were incubated with NAO to identify differences in membrane structure.

## Discussion

The LiaFSR envelope-stress-response pathway contributes to DAP resistance in both *E. faecalis* and *E. faecium*. When the response regulator of that system, LiaR, is deleted from the *E. faecium* genome, the isolate reverts to DAP hypersusceptibility (11). Therefore, a small molecule inhibitor of the LiaFSR pathway may extend DAP efficacy in the clinic. When *liaR* was deleted from the *E. faecium* genome, cells took approximately two weeks longer to evolve DAP resistance, regardless of environment (Figure 1). This delay supports the hypothesis that developing an inhibitor targeting LiaFSR, or LiaR specifically, could extend DAP efficacy against *E. faecium* and that alternative evolutionary trajectories did not provide rapid adaptation as might have been expected.

While resistance was significantly delayed in the absence of LiaR, evolution still provided a means to identify novel paths to DAP resistance. Here, the environment, once again (12), influenced how *E. faecium* evolved DAP resistance, independently of a functional LiaFSR system. Flask-transfer isolates broadly employed changes to the cell surface charge to mediate repulsion of DAP from the cell surface (Figure 2). This phenomenon was observed in both 503FΔ*liaR* and 515FΔ*liaR* isolates containing mutations to the *yvcRS* system which likely upregulated the production of *dltABCD* transcripts. While the sampling strategy did not provide a quantitative survey of all mutations derived from flask-adaptation, the concurrent evolution of *yvcRS* mutations across different populations and ancestral genomes suggests their importance in contributing to DAP resistance in this environment. Importantly, *yvcRS* mutations allowed *E. faecium* HOU503 to readily achieve high levels of DAP resistance (12); however, when *liaR* was deleted, this same level of resistance was not observed for an additional two weeks. This finding also supports previous work where a *yvcR* mutation was present in a DAP-resistant isolate of *E. faecalis* lacking *liaR* (34)

Interestingly, *yvcR*^G15S^ was observed in four 515FΔ*liaR* bioreactor-derived end-point isolates, yet they did not exhibit the repulsion phenotype (Table 4 and isolates P51/P53 in Figure 6). This may be due to the location of the mutation (N-terminus) compared to all other observed mutations that do show repulsion (C-terminus). Alternatively, the lack of repulsion may be due to the complexity of additional mutations present within these genomes.

While flask-transfer isolates predominately evolved resistance via repulsion, bioreactor-derived isolates evolved complex mechanisms of resistance that were heavily influenced by the ancestral genome. 503FΔ*liaR* bioreactor-derived isolates tended to display modest increases of cell surface charge, as observed in *divIVA*^I92F^ (Figure 4) and *murAA*^A149E^ (Figure 5). Alternatively, 515FΔ*liaR* bioreactor-derived isolates almost exclusively evolved one of two mutations in *metB* which resulted in diverse DAP adaptive phenotypes (Figures 6-7).

For isolates with mutations in *murAA*, it is possible that this mutation alters peptidoglycan synthesis leading to a remodeling of the cell wall, though there was no evidence of changes in cell wall thickness (Figure S5). It is possible that changes to the cell wall architecture may have decreased DAP access to the membrane, leading to increased resistance and are currently under investigation. These mutations, however, were observed in isolates with a variety of DAP MICs (ranging from 2-8 mg/L), suggesting that *murAA*^A149E^ alone was insufficient to confer resistance and additional mutations were required for high resistance levels.

In bioreactor-evolved 515FΔ*liaR* isolates, the presence of *metB* mutation alone did not appear to confer high levels of DAP resistance, as an isolate with the fewest additional mutations, isolate P24 (*metB*^+C^, sortase insertion^L8*^, *purK*^R153R^, *entf515F_691*^Δ155^), displayed a DAP MIC of 1 µg/ml. Additionally, this isolate did not display an increase in DAP-tolerance as assessed by growth on BHI agar in the presence of added glucose (Figure S7). In fact, there appears to be a strong relationship between secondary mutations that affect lipid metabolism and higher levels of DAP resistance. The most closely related end-point isolates to P24, P62 and P40 (*metB*^+C^, sortase insertion^L10*^, *purK*^R153R^, *fabI*^−108^) had DAP MICs of 4 and 8 µg/ml respectively, suggesting that the presence of a SNP upstream of *fabI*, which catalyzes the final step of fatty acid synthesis (36), contributed to higher levels of DAP resistance. Those isolates from the *metB*^+C^ lineage with the highest levels of DAP resistance (P29, P36, and P69) contained the sortase insertion^L10*^, *gdpD*^H29R^, and *entf515F_1865*^A327E^. GdpD plays a role in phospholipid metabolism and has been found in DAP-resistant isolates of *E. faecalis* and *E. faecium* (10, 12, 32), while *entf515F_1865* is annotated as a phospholipase family protein. These mutations support the hypothesis that lipid modification largely contributed to the DAP resistance phenotype and that the initial mutation in *metB* may have facilitated the success of these DAP resistant trajectories.

Membrane modifications remain important indicators of DAP resistance-specifically mutations to *cls.* While not directly conferring resistance, these mutations are found very frequently in DAP resistance populations of both *E. faecium* and *E. faecalis*, regardless of the underlying resistance mechanism or the presence of LiaFSR.

In summary, deleting *liaR* from the *E. faecium* genome greatly delayed the onset of DAP resistance. The evolved resistance mechanisms vary greatly depending on environment and ancestral genome, suggesting that deleting *liaR* positions *E. faecium* at a significant disadvantage and the organism struggled to find suitable resistance strategies. Because of this delay, it would be interesting to investigate whether the inhibition of LiaFSR signaling could be a useful strategy to extend DAP efficacy in the clinic.

## Materials and methods

### Bacterial strains and growth conditions

Clinical isolates *E. faecium* HOU503 and HOU515 (13) with deletions of *liaR* encoding the response regulator of the LiaFSR system were used (denoted as 503FΔ*liaR* and 515FΔ*liaR*, respectively) (11). Initial DAP MICs in Brain Heart Infusion (BHI) were 0.25 and 0.5 mg/L, respectively. All isolates were grown in BHI with added calcium (50 mg/L) unless otherwise stated. Additionally, *E. faecalis* S613 and R712 were used as controls in some experiments (35).

### Evolving DAP resistance in different environments by experimental evolution

Five populations of 503FΔ*liaR* and 515FΔ*liaR* each were adapted to DAP resistance using daily 100-fold dilutions via serial-flask-transfer as described previously (12). 503FΔ*liaR* and 515FΔ*liaR* were evolved in the bioreactor in singlicate as described previously (12, 17–21). Further details are found in the Supplemental Text.

### Genomic analysis of DAP-resistant isolates and populations

Whole genome sequencing (WGS) was performed on end-point isolates derived from the final populations. Deep sequencing was performed on the heterogenous daily bioreactor populations. Sequencing was performed with 2×150 bp reads on Hi-seq at Genewiz. Analysis was performed using the Breseq pipeline (36). Further details on sample collection, preparation and analysis are found in the Supplemental Text.

### Microscopic analysis of DAP-resistant isolates

End-point isolates were incubated with Poly-L-Lysine conjugated to the fluorophore FITC (PLL:FITC), DAP conjugated to the fluorophore bodipy (BDP:DAP), or 10-N-nonyl acridine orange (NAO) to determine DAP-resistant phenotypes as described previously (12) and more extensively in the Supplemental Text. Transmission Electron Microscopy (TEM) was performed at The University of Texas Health Science Center Electron Microscopy Core on a JEOL JEM 1200 EX Electron Microscope as described previously (12).

### RT-qPCR

RNA was extracted using the Qiagen RNeasy Mini kit with modifications (12) followed by Invitrogen DNAse I treatment and cDNA synthesis using Invitrogen SuperScript III in accordance with manufacturer’s instructions. Bio-Rad Sybr Green was used for qPCR on a Bio-Rad CFX Connect Real-Time System. Transcripts were compared to the house-keeping gene, glucose-1-dehydrogenase 4 (*gdhIV*), and are quantified using the 2^−ΔΔ*Ct*^ method. Experiments were performed in biological and technical triplicate.

### Statistical analysis

Statistical significance is defined as (p<0.05) using the Mann-Whitney test with *post hoc* Holm-Bonferroni adjustment unless otherwise stated.

### Data Availability

All sequences are deposited under PRJNA549910 (https://www.ncbi.nlm.nih.gov/bioproject/?term=PRJNA549910&utm_source=gquery&utm_medium=search).

## Transparency Declarations

C.A.A. has received grants from Merck, MeMEd Diagnostics, and Entasis Therapeutics. W.R.M. has received a grant from Merck, and honoraria from Achaogen and Shionogi. T.T. has received a grant from Merck.

